# Differential effects of PDE4A5 on cAMP-dependent forms of long-term potentiation

**DOI:** 10.1101/2024.05.04.592525

**Authors:** Satya Murthy Tadinada, Emily N. Walsh, Utsav Mukherjee, Ted Abel

## Abstract

cAMP signaling is critical for memory consolidation and certain of forms long-term potentiation (LTP). Phosphodiesterases (PDEs), enzymes that degrade the second messenger cAMP and cGMP, are highly conserved during evolution and represent a unique set of drug targets, given the involvement of these enzymes in several pathophysiological states including brain disorders. The PDE4 family of cAMP selective PDEs, exert regulatory roles in memory and synaptic plasticity, but the specific roles of distinct PDE4 isoforms in these processes are poorly understood. Building on our previous work demonstrating that spatial and contextual memory deficits were caused by expressing selectively the long isoform of the PDE4A subfamily, PDE4A5, in hippocampal excitatory neurons, we now investigate the effects of PDE4A isoforms on different cAMP-dependent forms of LTP. We find that PDE4A5 impairs long-lasting LTP induced by theta burst stimulation (TBS) while sparing long-lasting LTP induced by spaced 4-train stimulation (4X100Hz). This effect requires the unique N-terminus of PDE4A5 and is specific to this long isoform. Targeted overexpression of PDE4A5 in area CA1 is sufficient to impair TBS-LTP, suggesting that cAMP levels in the postsynaptic neuron are critical for TBS-LTP. Our results shed light on the inherent differences among the PDE4A subfamily isoforms, emphasizing the importance of the long isoforms, which have a unique N-terminal region. Advancing our understanding of the function of specific PDE isoforms will pave the way for developing isoform-selective approaches to treat the cognitive deficits that are debilitating aspects of psychiatric, neurodevelopmental, and neurodegenerative disorders.

**Key Points:**

- Hippocampal overexpression of a PDE4A subfamily long isoform, PDE4A5, but not a short isoform PDE4A1, impairs spatial and contextual fear memory and the N-terminus of PDE4A5 is important for this effect.
- Hippocampal overexpression of PDE4A isoforms, PDE4A1 and PDE4A5 do not impair LTP induced by spaced tetanic stimulation at the CA3-CA1 synapses.
- Hippocampal overexpression of PDE4A5, but not PDE4A1 or the N-terminus truncated PDE4A5 (PDE4A5Δ4) selectively impairs LTP induced by theta burst stimulation (TBS) at the CA3-CA1 synapses and expression of PDE4A5 in area CA1 is sufficient for the TBS-LTP deficit.
- These results suggest that PDE4A5, through its N-terminus, regulates cAMP pools that are critical for memory consolidation and expression of TBS-LTP at the CA3-CA1 synapses.

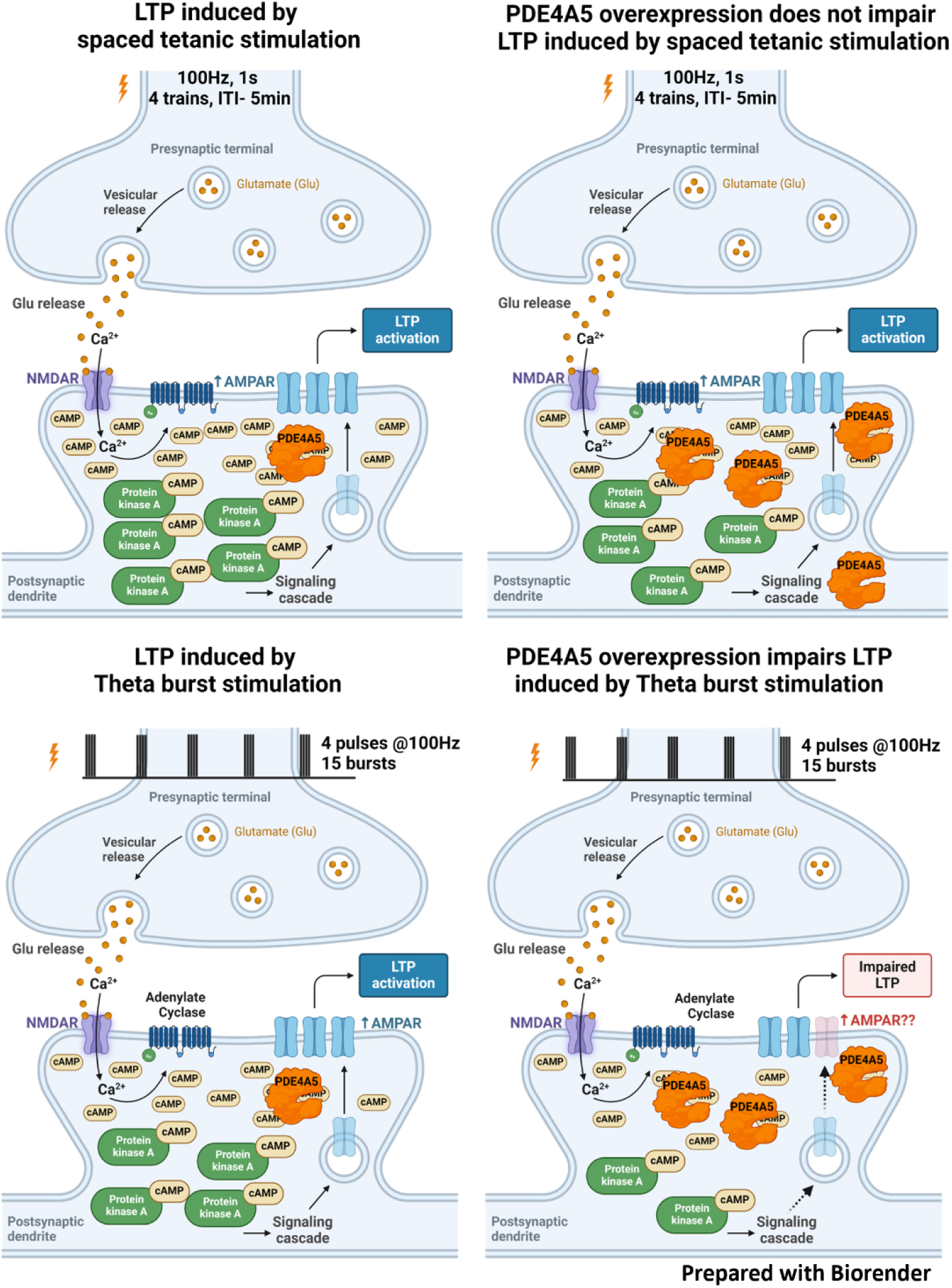
GRAPHICAL ABSTRACT. Spaced tetanic stimulation and TBS induce cAMP synthesis and activation of PKA to promote signaling cascades that facilitate expression of long-lasting LTP at the CA3-CA1 synapses. PDE4A5 overexpression in the hippocampus selectively impairs cAMP and PKA dependent TBS-LTP at the CA3-CA1 synapses, while sparing LTP induced by spaced tetanization.

## Introduction

Decades of research have established that the second messenger cAMP and downstream signaling pathways have an indispensable role in long-term memory consolidation and certain long-lasting forms of synaptic plasticity (Frey *et al*., 1993; Abel *et al*., 1997; Wong *et al*., 1999; Ma *et al*., 2009; Kandel, 2012). Protein kinase A (PKA), exchange proteins activated by cAMP (Epacs), Popeye domain containing proteins, and cyclic nucleotide-gated ion channels (CNG) mediate the downstream effects of cAMP on cellular function and physiology. Modulation of intracellular signaling by dynamic changes in cAMP depends on the spatial and temporal regulation of cAMP levels (Bock *et al*., 2020; Zhang *et al*., 2020; Zaccolo *et al*., 2021), and cAMP degradation by phosphodiesterases is an evolutionarily conserved mechanism for regulation of intracellular cAMP levels. The mammalian genome contains a total of 11 PDE families (PDE 1-11) with cAMP-selective phosphodiesterases belonging to the PDE 4, 7, and 8 families, cGMP-selective phosphodiesterases belonging to the PDE 5, 6, and 9 families and phosphodiesterases belonging to PDE 1, 2, 3, 10 and 11 families lack selectivity and degrade both cAMP and cGMP (Keravis & Lugnier, 2012). The PDE4 family of phosphodiesterases accounts for a significant proportion of cAMP phosphodiesterase activity in the brain, and the importance of PDE4 activity in the brain is also reflected in the efforts to develop PDE4 inhibitors as cognitive enhancers to treat Alzheimer’s disease (Richter *et al*., 2013; Tibbo *et al*., 2019) and Fragile X syndrome (Berry-Kravis *et al*., 2021).

The significance of cAMP and PKA in behavioral sensitization was identified in Aplysia (Brunelli *et al*., 1976; Castellucci *et al*., 1980), while the mechanisms that regulate intracellular cAMP levels for learning and memory was highlighted by findings in *Drosophila* (Dudai *et al*., 1976; Byers *et al*., 1981; Livingstone *et al*., 1984). Mutations in either *rutabaga* (Livingstone *et al*., 1984) or in *dunce* (Dudai *et al*., 1976; Byers *et al*., 1981), which resulted in a decrease and an increase in cAMP respectively, caused memory deficits in aversive odor learning tasks. Similar observations were reported in *Aplysia*, where overexpression or knockdown of apPDE resulted in deficits in synaptic facilitation (Park *et al*., 2005). In mice, concomitant loss of the Ca^+2^ sensitive adenylyl cyclases, AC1 and AC8, resulted in memory and LTP deficits (Wong *et al*., 1999). Administration of rolipram, a pan-PDE4 inhibitor, enhances fear conditioning memory and LTP at low doses (Barad *et al*., 1998; Navakkode *et al*., 2004), suggesting that phosphodiesterases may function as a memory suppressor gene (Abel *et al*., 1998), serving as a brake on synaptic potentiation and memory consolidation.

The mammalian PDE4 family of phosphodiesterases consists of 4 genes that encode the subfamilies PDE4A-D, giving rise to at least 25 transcripts generated from multiple transcription start sites as well as alternative splicing (Paes *et al*., 2021). PDE4A, 4B, and 4D subfamily isoforms primarily contribute to the overall PDE4 activity in the brain, and studies in mutant mouse lines have provided fundamental insights into the function of these various PDE4 subfamilies. Knockout of the mammalian homolog of *dunce,* PDE4D, confers resilience to depression (Zhang *et al*., 2002), enhances adult neurogenesis and spatial memory (Li *et al*., 2011), but impairs fear memory (Rutten *et al*., 2008). On the other hand, the knockout of PDE4A (Hansen *et al*., 2014) or PDE4B (Zhang *et al*., 2008) results in increased anxiety-like phenotypes. Although these studies shed light on our understanding of the role of individual PDE4 subfamilies in various behaviors, the fact that these models are global knockouts means that all isoforms within a subfamily are removed in these mutant mice, limiting our ability to identify the function of individual isoforms within each subfamily. As demonstrated in our earlier work (Havekes *et al*., 2016a) this limitation can be circumvented by selectively overexpressing specific PDE4 isoforms in cell types in the brain to investigate the effects on cognition. Overexpression of the long isoform of the PDE4A subfamily, PDE4A5, but not the super-short isoform, PDE4A1, in excitatory neurons in the hippocampus, selectively impairs long-term spatial and contextual memory without inducing anxiety like phenotypes (Havekes *et al*., 2016a).

To further our understanding of the role of the PDE4A subfamily in synaptic plasticity, we investigated the effects of overexpression of PDE4A subfamily isoforms on LTP. We focused on PDE4A1 and PDE4A5 because these two isoforms are the most abundant isoforms at the protein level in the hippocampus (Cherry & Davis, 1999; Hansen *et al*., 2014). We find that PDE4A5 overexpression differentially affects cAMP-dependent forms of long-lasting plasticity such that long-lasting LTP induced by Spaced 4-train stimulation (4X100Hz) is spared, while long-lasting LTP induced by burst stimulation in theta frequency (TBS) is selectively impaired. We further provide evidence that the TBS-LTP deficit is selective to the long isoform PDE4A5 and requires the unique N-terminus of PDE4A5. Our results suggest that postsynaptic expression of PDE4A5 is sufficient for the deficit in TBS-LTP and that subcellular localization of PDE4A5 may, in part, contribute to these effects.

## Methods

### Animals

All procedures were approved by the Institutional Animal Care and Use Committee at The University of Iowa. C57BL/6J mice were purchased from The Jackson Laboratories (Catalog# 000664, RRID: IMSR_JAX:000664, Bar Harbor, ME) and were housed in continuously ventilated standard caging with access to food and water ad libitum. Housing rooms were maintained on a 12h light: dark cycle and under constant conditions for humidity and temperature.

### Surgeries

8 to 12-week-old animals were subject to stereotaxic surgeries, and Adeno-associated viruses (AAVs) used in the study were previously described (Havekes *et al*., 2016a). AAV2/9 particles were bilaterally injected to express PDE4A isoforms or GFP (Addgene #50469) in the hippocampal excitatory neurons using the CaMKII promoter, and titers of the various AAVs used in this study were adjusted to 1E+13 genome copies per mL. 1µL was delivered into each hemisphere (stereotaxic coordinates: A/P-1.9mm from bregma, M/L- ±1.5mm and D/V- 1.5mm from the surface of the brain) over 5 minutes. For CA1 selective expression, 350nL of the 1E+13 genome copies per mL titer was used, and stereotaxic coordinates were validated as A/P- 2.1mm from bregma, M/L-±1.25mm and D/V- 1.2mm from the surface of the brain. Animals were allowed 4-6 weeks of recovery following surgery. In all experiments, eGFP was the control group, and PDE4A isoform overexpression was the experimental group.

### Immunohistochemistry

Immunohistochemistry was performed as previously described (Havekes *et al*., 2016b). Infusion-fixed brains (4% PFA) were stored in 30% (w/v) sucrose solution at 4°C until further processing. 30µm coronal sections were prepared (Leica Cryostat CM3050S, Leica Biosystems, Wetzlar, Germany) and rinsed three times in 1X PBS for 10 minutes. After one hour incubation in blocking solution (3% goat serum and 0.2% TritonX-100 in 1X PBS) at room temperature, sections were incubated in primary antibody (1:200 dilution at 4°C for 48h) against the epitope tags (VSV-G tag for PDE4A5 at 1:200 dilution, ab1874, RRID: AB_302646, Abcam, Cambridge, UK; HA tag for PDE4A1 and PDE4A5Δ4 at 1:200 dilution, CS3724, RRID: AB_1549585, Cell signaling, Danvers, MA). Sections were rinsed three times in 1X PBS and incubated at room temperature for two hours with Alexa fluor 594 conjugated secondary antibodies (1:10,000 dilution; donkey anti-rabbit, A21207, RRID: AB_141637, Invitrogen, Waltham, MA). Sections were mounted on super frost plus slides (Catalog # 12-550-15, Fisher Scientific, Pittsburgh, PA) with medium containing the nuclear stain DAPI (P36962, Invitrogen, Waltham, MA). In experiments involving CA1 targeted expression of PDE4A5, post-recording immunohistochemistry for VSV was performed and used as inclusion criteria (see Figure 5H, H1-H4). Slices used in LTP recordings from animals that received AAV for PDE4A5 overexpression, but lacking VSV signal in CA1 as determined by IHC post-hoc were categorized as PDE4A5 negative (PDE4A5 -ve).

### cAMP-Phosphodiesterase assay

cAMP-PDE assay (ab273342, Abcam, Cambridge, MA) was performed according to the protocol provided by the manufacturer with minor modifications. Reactions were scaled down by proportionately matching the components to a final volume of 20uL and run in 384-well format instead of 96 wells. Samples were run in duplicates, and background subtracted average values were used to determine PDE activity from a standard curve generated using the AMP standard provided by the manufacturer. Fluorescence measurements (Ex/Em 535nm/587nm) were made for 30 minutes on a Cytation 5 plate reader (BioTek, Winooski, VT) with PMT gain set at 50 and a read interval of 2 minutes.

### Slice Preparation

Transverse hippocampal slices were prepared as described previously (Shetty *et al*., 2015). Briefly, animals were sacrificed by cervical dislocation followed by decapitation, and the brain was quickly removed and placed in ice-cold aCSF (124 mM NaCl, 4.4 mM KCl, 2.5 mM CaCl_2_, 1.3 mM MgCl_2_, 1 mM NaH_2_PO_4_, 10 mM glucose and buffered with 26.2 mM NaHCO_3_) bubbled constantly with carbogen and the hippocampus was dissected out with a sickle scaler. 350-400µm thick transverse slices were then prepared using a McIlwain tissue chopper (Stoelting, Wood Dale, IL), and slices were immediately transferred to a nylon net insert on a humidified interface chamber constantly perfused with aCSF (saturated with carbogen) at a flow rate of 1mL/min and were allowed to recover at 30°C for at least two hours before recordings were performed.

### Electrophysiology

Extracellular field recordings were performed by stimulation of Schaffer-collaterals using a monopolar stimulation electrode fabricated from nichrome wire (Catalog # 762000, A-M systems, Sequim, WA). Both stimulation and recording electrodes were placed within area CA1 and pulled glass electrodes (Catalog # B150-110-10, Sutter Instruments, Novato, CA) filled with aCSF (2-5MΩ resistance) were used for recording. An isolated pulse stimulator (Model 2100, A-M systems, Sequim, WA) was used to inject constant current pulses at increasing intensity (5-50µA) to generate an input-output curve and stimulation intensity (test stimulation) corresponding to 30-40% of maximum fEPSP amplitude was used for baseline recordings. Paired pulse facilitation was assessed by delivering a pair of pulses at test stimulation intensity (30-40% of maximum fEPSP amplitude) with an interval ranging between 25-300ms. After a 20-minute baseline recording, LTP induction was performed at the same stimulation intensity as test stimulation using one of 3 paradigms-spaced 4-train stimulation, massed 4-train stimulation, or theta-burst stimulation. Spaced 4-train and massed 4-train induction involved delivering four 1s 100Hz trains at test stimulation intensity repeated with an inter-trial interval of 5 minutes (spaced) or 5 seconds (massed). Theta burst LTP was induced by delivering 15 bursts of 4 pulses, with pulses delivered at 100Hz and bursts delivered at theta frequency (5Hz, interburst interval-200ms). Signals were amplified using an intracellular electrometer (IE251A, Warner Instruments, Holliston, MA) and low-pass filtered at 2kHz using a low-pass Bessel filter (LPF-100B, Warner Instruments, Holliston, MA). The sampling frequency was set at 20kHz, and signals were digitized using an Axon Digidata 1440A digitizer (Axon Instruments, Molecular Devices, San Jose, CA) controlled by pClamp 11 software. Custom protocols written using pClamp v10 and v11 used the digitizer output function to deliver a TTL pulse to the stimulator to deliver the analog signals. Data analysis was performed offline to measure the fEPSP slopes (20% to 80%), and slopes of post-LTP induction recordings are presented as % relative to the average slope of the baseline period, which was set to 100%.

### Statistics

Statistical analysis was performed using GraphPad Prism v9 or later to determine significant differences between groups with significance set at *P*<0.05. Repeated measures Two-way ANOVA was chosen for repeated measures data, and Welch’s One-way ANOVA was performed when more than two groups were compared and equal variances could not be assumed. Post-hoc analysis using Dunnett’s T3 multiple comparison test was used to determine significant differences. For comparison between two groups, unpaired Welch’s t-test was used. *N* reflects the number of animals in all the experiments. Data are presented as Mean ± SD.

## Results

### PDE4A isoform overexpression increases hippocampal cAMP PDE activity

To investigate the effects of overexpression of PDE4A isoforms on hippocampal phosphodiesterase activity, we bilaterally injected AAVs to express eGFP, PDE4A1, PDE4A5, or the N-terminal truncated PDE4A5Δ4 in excitatory cells of the hippocampus of wildtype mice. The forms of PDE4A chosen were based on the domain structure of this subfamily (Figure 1A) and their abundance in the hippocampus (Cherry & Davis, 1999; Hansen *et al*., 2014). All isoforms within the PDE4A subfamily contain a C-terminal region that includes the catalytic domain and the subfamily-specific C-terminus. The long isoform, PDE4A5, has a unique N-terminus followed by upstream conserved region 1 (UCR1) and upstream conserved region 2 (UCR2) that leads into the C-terminal region. The N-terminus of the super-short form, PDE4A1, forms α-helices and is required for membrane insertion of this isoform (Shakur *et al*., 1993; Scotland & Houslay, 1995; Shakur *et al*., 1995). PDE4A1 lacks UCR1 and has a truncated upstream conserved region (UCR2), followed by the remainder of the C-terminus. PDE4A5Δ4 is an experimental construct that lacks the N-terminus of PDE4A5; the region suggested to confer unique properties through protein-protein interactions and differential localization in various subcellular domains (Beard *et al*., 2002).

**Figure 1.**
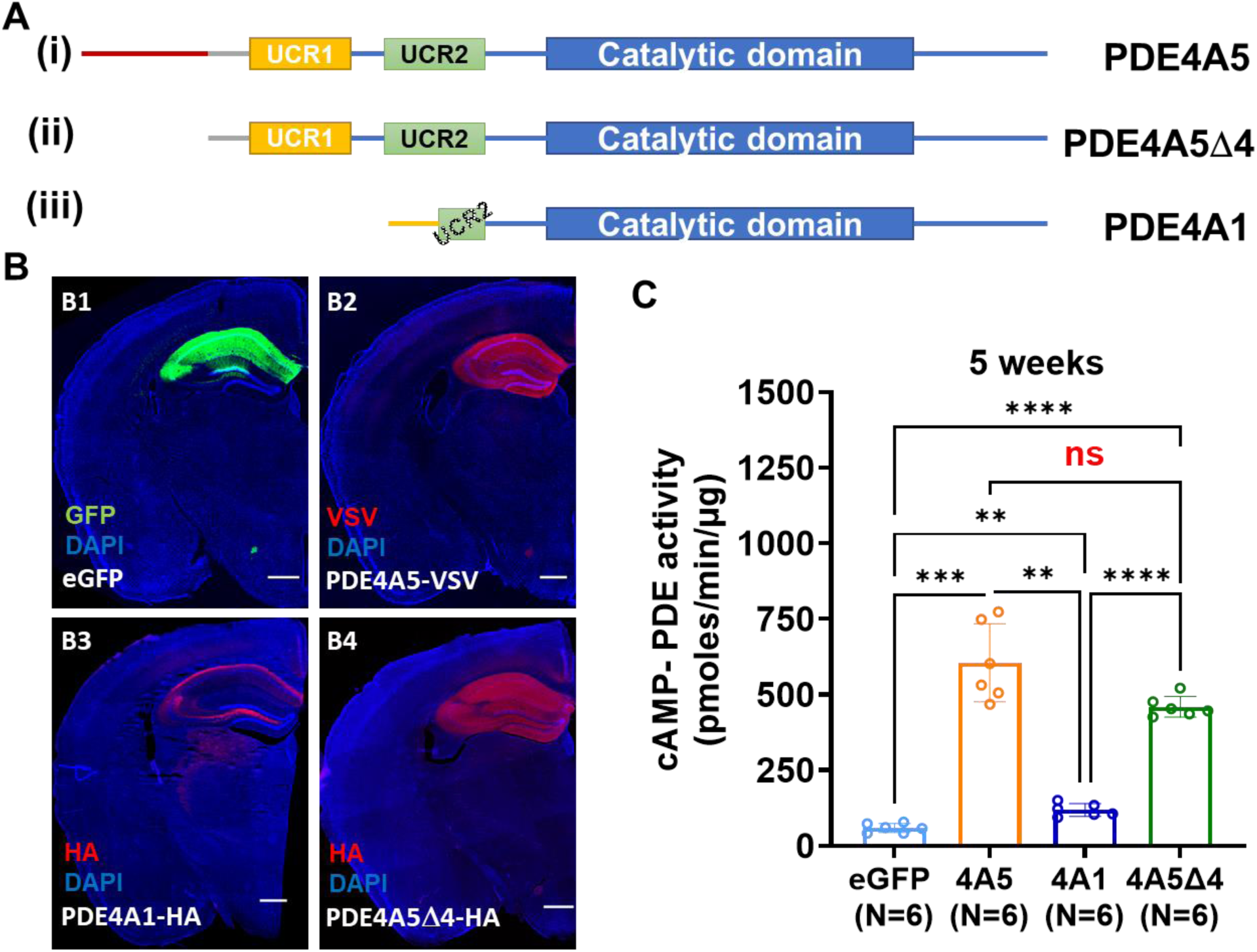
Validation of PDE4A overexpression and activity in the hippocampus. (A) Domain structure of PDE4A subfamily isoforms investigated in this study. The red line in (i) represents the unique N-terminus of PDE4A5, and the yellow line in (iii) represents the unique N-terminus of PDE4A1. (B) Immunohistochemistry showing expression of (B1) eGFP or (B2-B4) PDE4A isoforms in the hippocampus of mice four weeks post-surgery. Primary antibodies against (B2) VSV and (B3, B4) HA epitopes showed expression of (B2) PDE4A5, (B3) PDE4A1, and (B4) PDE4A5Δ4 respectively, restricted primarily to the hippocampus (C) cAMP-PDE activity in the hippocampus of mice overexpressing PDE4A isoforms at five weeks post-surgery (Mean ± SD of cAMP-PDE activity in pmol/min/µg, eGFP: 58.95 ± 15.18, PDE4A5: 605.2 ± 128.9, PDE4A1: 118.3 ± 21.3, PDE4A5Δ4: 460.0 ± 34.5; *P* < 0.0001 by Brown-Forsythe ANOVA test followed by Dunnett’s T3 multiple comparison tests, F* (3, 6.154) = 90.6; *P*=0.0007 for eGFP vs. PDE4A5, *P* = 0.0019 for eGFP vs. PDE4A1, *P* = 0.0013 for PDE4A5 vs. PDE4A1, *P* = 0.1638 for PDE4A5 vs. PDE4A5Δ4, *P*<0.0001 for eGFP vs. PDE4A5Δ4 and PDE4A1 vs. PDE4A5Δ4)

Viral expression of PDE4A isoforms was seen in pyramidal neurons throughout the hippocampus (Figure 1B). Unlike PDE4A5 and PDE4A5Δ4, which showed labeling throughout the hippocampal fields (Figures B2 and B4), PDE4A1 showed selectively high levels of expression in the alveus and stratum oriens with little to no expression in stratum radiatum (Figure B3). Overexpression of PDE4A constructs increased hippocampal cAMP selective phosphodiesterase activity by 10-fold when PDE4A5 was overexpressed and up to 8-fold when PDE4A5Δ4 was overexpressed. Overexpression of PDE4A1 resulted in a 2-fold increase in overall cAMP-PDE activity (Mean ± SD of cAMP-PDE activity in pmol/min/µg, eGFP: 58.95 ± 15.18, PDE4A5: 605.2 ± 128.9, PDE4A1: 118.3 ± 21.3, PDE4A5Δ4: 460.0 ± 34.5; *P* < 0.0001 by Brown-Forsythe ANOVA test followed by Dunnett’s T3 multiple comparison tests, F* (3, 6.154) = 90.6; *P*=0.0007 for eGFP vs. PDE4A5, *P* = 0.0019 for eGFP vs. PDE4A1, *P* = 0.0013 for PDE4A5 vs. PDE4A1, *P* = 0.1638 for PDE4A5 vs. PDE4A5Δ4, *P*<0.0001 for eGFP vs. PDE4A5Δ4 and PDE4A1 vs. PDE4A5Δ4). The relatively smaller increase in cAMP phosphodiesterase activity upon PDE4A1 overexpression may be related to the localization of this isoform selectively in the alveus and stratum oriens, and could also be due to potential limitations in the solubility of this isoform in the assay buffer because of the near complete association of PDE4A1 with the membranes (Shakur *et al*., 1993). *PDE4A isoform overexpression does not alter basal synaptic transmission*.

To investigate the effects of the overexpression of PDE4A isoforms on hippocampal synaptic plasticity, acute hippocampal slices were prepared from the brains of animals overexpressing eGFP or PDE4A isoforms 4-6 weeks after surgery. We first assessed the impact of PDE4A isoform overexpression on basal synaptic transmission properties by examining the input-output relationship and paired-pulse facilitation (PPF). Input-Output relationship showed increases in fEPSP amplitudes and presynaptic fiber volleys (PFV) in response to increasing stimulation intensities in all the groups tested (Figure 2A-C). Comparison of input-output relationship in slices from animals overexpressing PDE4A isoforms (experimental groups) compared to eGFP (control group) revealed no significant differences between the groups (Figure 2A-C; for main effects of PDE isoform expression on PFV amplitude, F(3,77) = 0.6725, *P* = 0.5715 by repeated measures two-way ANOVA. For main effects of PDE isoform expression on fEPSP amplitude, F(3,77) = 1.973, *P* = 0.1250 by repeated measures two-way ANOVA). Paired-pulse facilitation, a form of short-term plasticity believed to be primarily presynaptic (Zucker & Regehr, 2002; Jackman & Regehr, 2017), was measured as a relative change in fEPSP amplitude elicited by the second pulse compared to the first pulse when the two pulses are delivered with a short intertrial interval (ranging from 25-300ms in our experiments). PDE4A isoform overexpression did not result in significant differences in paired-pulse facilitation compared to the control eGFP group (Figure 2D, for main effects of PDE isoform expression, F(3,28) = 1.095, *P* = 0.3630 by repeated measures two-way ANOVA). These data suggest that PDE4A isoform overexpression does not influence basal synaptic transmission or short-term plasticity.

**Figure 2.**
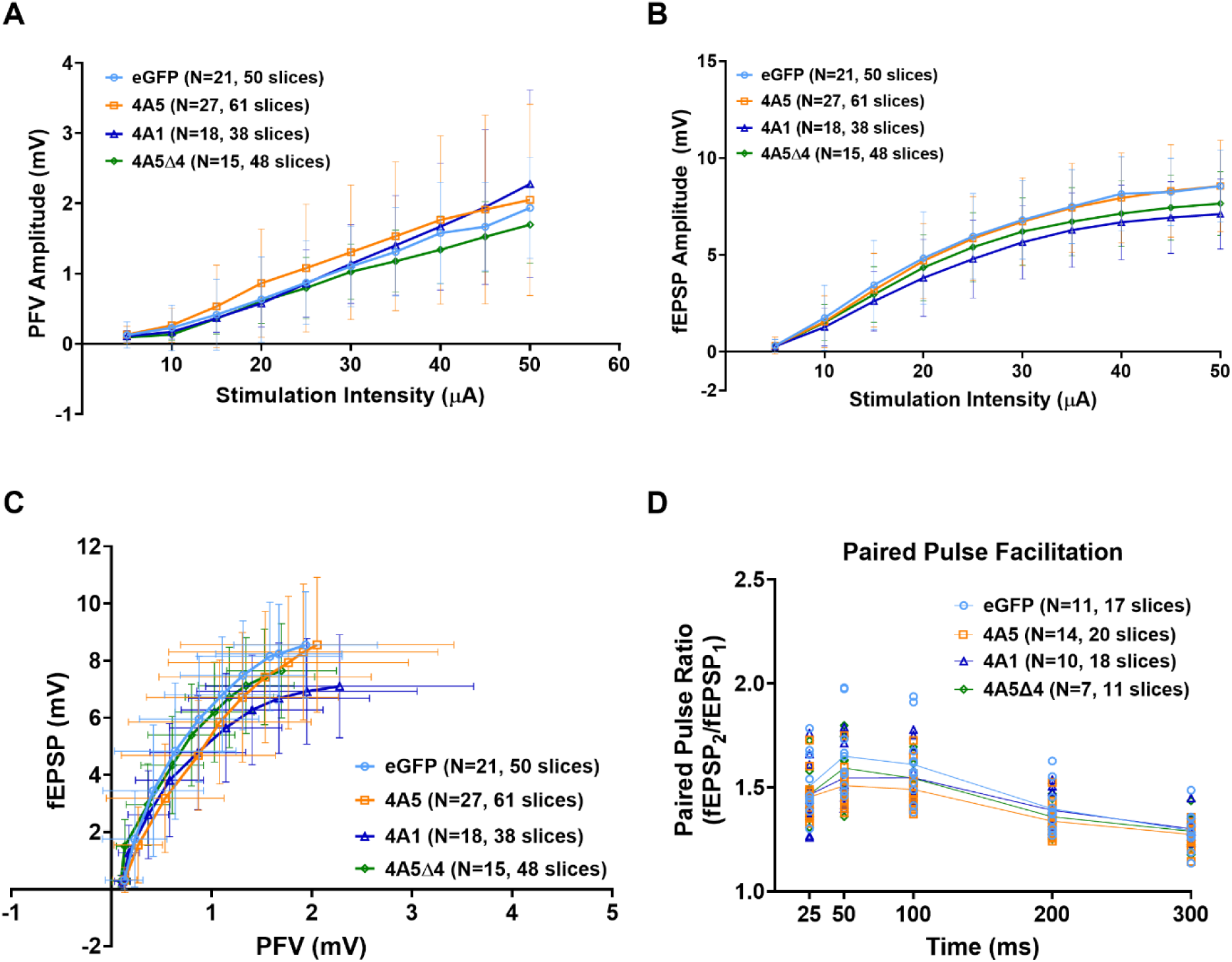
Overexpression of PDE4A isoforms does not affect basal synaptic transmission. Relationship between stimulation intensity and (A) Presynaptic Fiber Volley (PFV; for main effects related to PDE overexpression, F(3,77) = 0.6725, *P* = 0.5715 by repeated measures two-way ANOVA) (B) Field Excitatory Postsynaptic potentials (fEPSP) in slices overexpressing eGFP or PDE4A isoforms (for main effects related to PDE overexpression, F(3,77) = 1.973, *P* = 0.1250 *P* = 0.1250 by repeated measures two-way ANOVA for fEPSP amplitudes) (C) Input-Output relationship showing the relationship between presynaptic fiber volleys and field EPSPs in slices from animals expressing eGFP or overexpressing PDE4A isoforms (D) Paired Pulse facilitation showing facilitation of fEPSP in slices overexpressing eGFP and PDE4A isoforms (for main effects of PDE isoform expression, F(3,28) = 1.095, *P* = 0.3630 by repeated measures two-way ANOVA).

### PDE4A overexpression does not impair long-lasting plasticity induced by repeated tetanic stimulation

Next, we investigated the effects of PDE4A isoform overexpression on cAMP-dependent forms of long-lasting LTP in the CA1 region (Figure 3A). When LTP was induced by a spaced 4-train paradigm that elicits a cAMP/PKA dependent form of LTP (Abel *et al*., 1997; Woo *et al*., 2000, 2003), we found that overexpression of PDE4A5 and PDE4A5Δ4 resulted in a slightly lower post-tetanic potentiation measured 1 minute after the first tetanus, compared to slices from hippocampi overexpressing eGFP (Figure 3B; Mean ± SD relative to baseline- eGFP: 332.5 ± 61.49, PDE4A5: 262.1 ± 61.06, PDE4A1: 336.3 ± 87.97, PDE4A5Δ4: 246.8 ± 67.94, N ≥ 8 animals per group, *P* = 0.0144 by Welch’s ANOVA test, W(3,16.34) = 4.761; Dunnet’s multiple comparisons- eGFP vs. 4A5Δ4, *P* = 0.0385; *P* < 0.05 by Welch’s two-tailed t-test- eGFP vs. 4A5: *t* = 2.365, df = 14.79, *P* = 0.0321; eGFP vs. 4A5Δ4: *t* = 2.807, df = 15.84, *P* = 0.0128). However, there were no significant differences in the maintenance of LTP measured 2h after the last tetanus in slices overexpressing PDE4A isoforms compared to eGFP (Figure 3C, Mean ± SD of last 20 min, 100-120 min after the last tetanus relative to baseline- eGFP: 192.1 ± 66.47, PDE4A5: 197.5 ± 53.53, PDE4A1: 224 ± 45.97, PDE4A5Δ4: 198 ± 64.45, N ≥ 8 animals per group, *P* = 0.6211 by Welch’s ANOVA test, W(3,16.63) = 0.6046). These results suggest that PDE4A5 and PDE4A1 overexpression minimally affect LTP induced by spaced 4-train stimulation.

**Figure 3.**
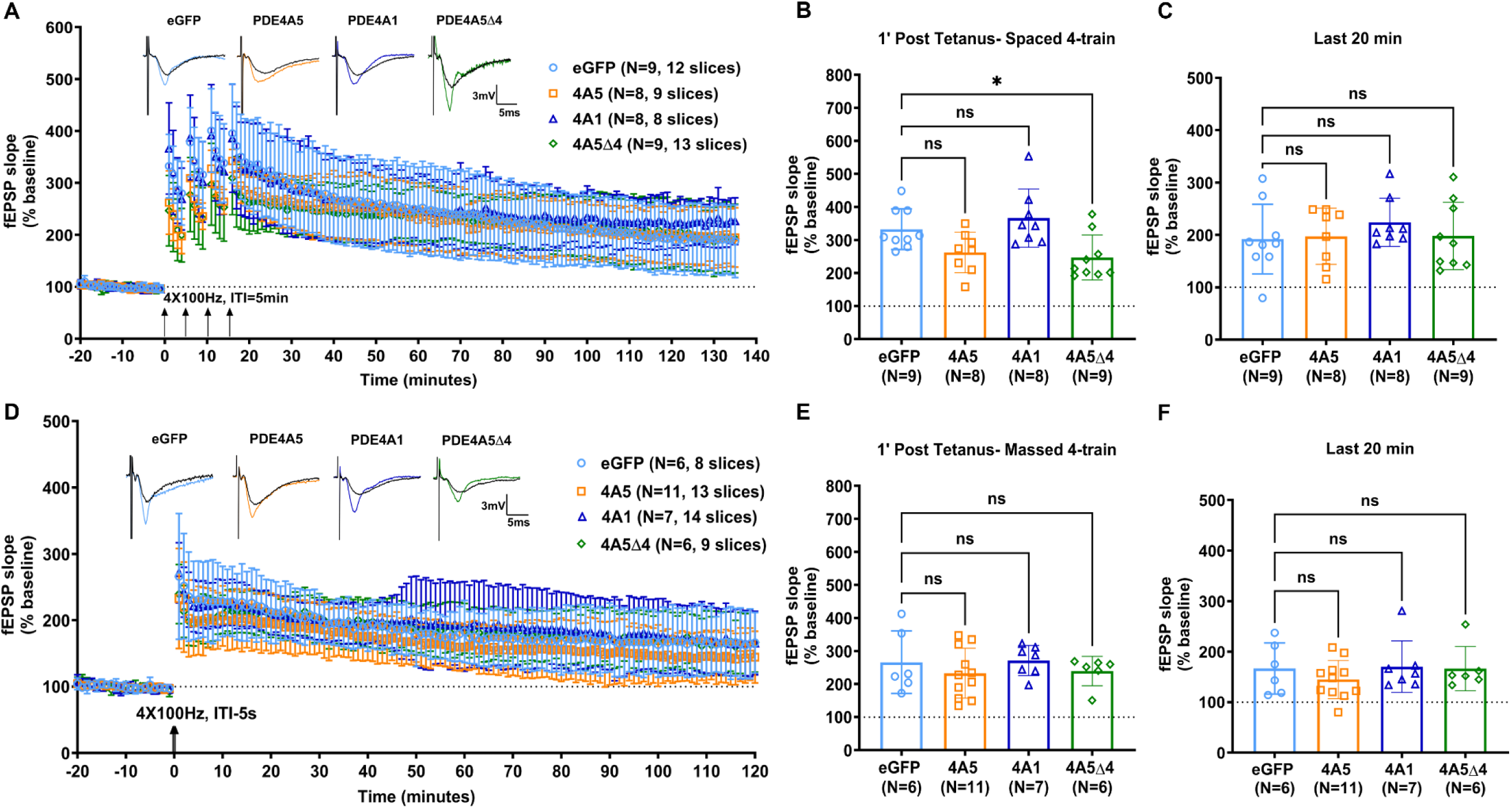
PDE4A overexpression does not impair cAMP-dependent and cAMP-independent forms of long-lasting LTP induced by repeated tetanic stimulation. (A) Overexpression of PDE4A isoforms did not affect spaced 4-train LTP. Representative traces shown at time zero (black trace) and at 120 min after the last tetanus (colored trace) for each group (B) fEPSP slopes 1 minute after the first 100Hz tetanus (Mean ± SD relative to baseline- eGFP: 332.5 ± 61.49, PDE4A5: 262.1 ± 61.06, PDE4A1: 336.3 ± 87.97, PDE4A5Δ4: 246.8 ± 67.94, N ≥ 8 animals per group, *P* = 0.0144 by Welch’s ANOVA test, W(3,16.34) = 4.761; Dunnet’s multiple comparisons- eGFP vs. 4A5Δ4, *P* = 0.0385; *P* < 0.05 by Welch’s two- tailed t-test- eGFP vs. 4A5: *t* = 2.365, df = 14.79, *P* = 0.0321; eGFP vs. 4A5Δ4: *t* = 2.807, df = 15.84, *P* = 0.0128) (C) maintenance of L-LTP induced by Spaced 4-train stimulation (Mean ± SD of last 20 min, 100-120 min after the last tetanus relative to baseline- eGFP: 192.1 ± 66.47, PDE4A5: 197.5 ± 53.53, PDE4A1: 224 ± 45.97, PDE4A5Δ4: 198 ± 64.45, N ≥ 8 animals per group, *P* = 0.6211 by Welch’s ANOVA test, W(3,16.63) = 0.6046). (D) Overexpression of PDE4A isoforms did not affect long-lasting LTP induced by massed 4-train stimulation. Representative traces shown at time zero (black trace) and at 120 min post-LTP induction (colored trace) for each group (E) fEPSP slopes 1 minute after the massed 4-train stimulation (Mean ± SD relative to baseline: eGFP: 266.1 ± 94.78, PDE4A5: 232.7 ± 75.48, PDE4A1: 271.4 ± 45.53, PDE4A5Δ4: 239.3 ± 44.6, N ≥ 6 animals per group, *P* = 0.6099 by Welch’s ANOVA test, W(3, 12.94) = 0.7867) (F) maintenance of LTP induced by massed 4-train stimulation (Mean ± SD of last 20 min, 100-120 min after tetanic stimulation relative to baseline- eGFP: 166.7 ± 50.55, PDE4A5: 144.8 ± 37.84, PDE4A1: 170.1 ± 50.99, PDE4A5Δ4: 166.5 ± 43.82, N ≥ 6 animals per group, *P* = 0.6078 by Welch’s ANOVA test, W(3, 12.04) = 0.6328).

We also investigated the impact of PDE4A isoform overexpression on massed 4-train stimulation-induced LTP, a form of LTP that has been demonstrated to be independent of cAMP and PKA (Woo *et al*., 2003) and find that massed 4-train induced LTP is unaffected by hippocampal overexpression of PDE4A isoforms (Figure 3D). Post-tetanic potentiation measured 1 minute after massed 4-train stimulation was not affected in slices with exogenous PDE4A isoform expression compared to eGFP expression (Figure 3E, Mean ± SD relative to baseline: eGFP: 266.1 ± 94.78, PDE4A5: 232.7 ± 75.48, PDE4A1: 271.4 ± 45.53, PDE4A5Δ4: 239.3 ± 44.6, N ≥ 6 animals per group, *P* = 0.6099 by Welch’s ANOVA test, W(3, 12.94) = 0.7867). Similar to spaced 4-train LTP, maintenance of LTP in slices from animals overexpressing PDE4A isoforms was not significantly different compared to eGFP overexpression (Figure 3F; Mean ± SD of last 20 min, 100-120 min after tetanic stimulation relative to baseline- eGFP: 166.7 ± 50.55, PDE4A5: 144.8 ± 37.84, PDE4A1: 170.1 ± 50.99, PDE4A5Δ4: 166.5 ± 43.82, N ≥ 6 animals per group, *P* = 0.6078 by Welch’s ANOVA test, W(3, 12.04) = 0.6328) suggesting that PDE4A isoform overexpression does not interfere with the maintenance of LTP induced by repeated tetanic stimulation.

### PDE4A5 overexpression impairs long-lasting plasticity induced by theta burst stimulation

We then investigated the effect of PDE4A isoform overexpression on synaptic plasticity induced by theta burst stimulation, another cAMP/PKA signaling-dependent form of long-lasting LTP (Nguyen & Kandel, 1997). The stimulation pattern used in TBS reflects the firing patterns of neurons during exploratory behavior, and the plasticity resulting from TBS is believed to be induced by a more physiologically relevant firing pattern (Larson *et al*., 1986). Compared to slices from animals overexpressing eGFP, post-tetanic potentiation measured 1 min after TBS was lower in slices from animals overexpressing PDE4A5 alone, but not in slices from animals overexpressing the other isoforms (Figure 4B; Mean ± SD relative to baseline: eGFP: 250.9 ± 53.97, PDE4A5: 195.4 ± 40.39, PDE4A1: 284 ± 82.51, PDE4A5Δ4: 287.2 ± 80.55, N ≥ 8 animals per group, *P* = 0.0061 by Welch’s ANOVA test, W (3, 16.63) = 5.929; Dunnet’s multiple comparisons: *P* = 0.0765 for eGFP vs. PDE4A5; *P* < 0.05 by Welch’s two-tailed t-test: eGFP vs. 4A5: *t* = 2.51, df = 11.84, *P* = 0.0277). Importantly, LTP induced by this pattern of stimulation decayed back to baseline over time in slices overexpressing PDE4A5 as compared to slices from animals overexpressing eGFP or other PDE4A isoforms (Figure 4A). Thus, LTP maintenance was significantly impaired only by overexpression of PDE4A5 (Figure 4C, Mean ± SD of last 20 min, 100-120 min after TBS relative to baseline- eGFP: 165.7 ± 43.73, PDE4A5: 105 ± 19.81, PDE4A1: 156.4 ± 32.9, PDE4A5Δ4: 173.8 ± 50.73, N≥8 animals per group, *P* = 0.0003 by Welch’s ANOVA test, W(3,15.63) = 11.53; Dunnet’s multiple comparisons- *P* = 0.0141 for eGFP vs. PDE4A5). These results suggest that increased activity of PDE4A5, in hippocampal excitatory neurons is detrimental to long-lasting plasticity induced by theta burst stimulation.

**Figure 4.**
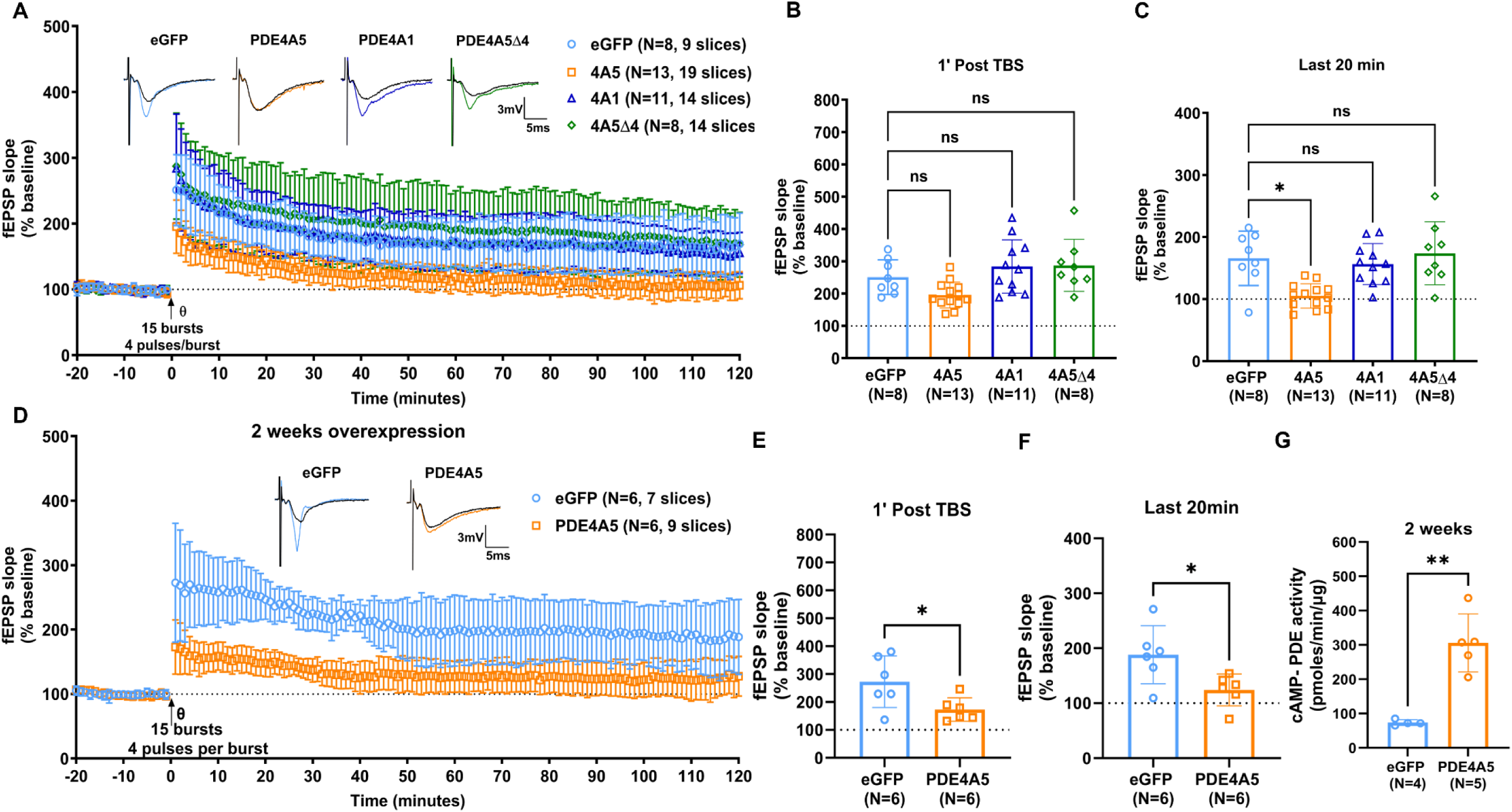
PDE4A5 overexpression impairs cAMP-dependent form of long-lasting LTP induced by theta burst stimulation. (A) LTP induced by TBS is impaired by PDE4A5 overexpression but not by PDE4A1 or PDE4A5Δ4 overexpression. Representative traces shown at time zero (black trace) and at 120 min post-TBS (colored trace) for each group (B) fEPSP slope 1 minute after theta burst stimulation (Mean ± SD relative to baseline: eGFP: 250.9 ± 53.97, PDE4A5: 195.4 ± 40.39, PDE4A1: 284 ± 82.51, PDE4A5Δ4: 287.2 ± 80.55, N ≥ 8 animals per group, *P* = 0.0061 by Welch’s ANOVA test, W (3, 16.63) = 5.929; Dunnet’s multiple comparisons: *P* = 0.0765 for eGFP vs. PDE4A5; *P* < 0.05 by Welch’s two-tailed t-test: eGFP vs. 4A5: *t* = 2.51, df = 11.84, *P* = 0.0277) (C) PDE4A5 overexpression resulted in a deficit in the maintenance of LTP induced by TBS (Mean ± SD of last 20 min, 100-120 min after TBS relative to baseline- eGFP: 165.7 ± 43.73, PDE4A5: 105 ± 19.81, PDE4A1: 156.4 ± 32.9, PDE4A5Δ4: 173.8 ± 50.73, N≥8 animals per group, *P* = 0.0003 by Welch’s ANOVA test, W(3,15.63) = 11.53; Dunnet’s multiple comparisons- *P* = 0.0141 for eGFP vs. PDE4A5) (D) TBS-LTP deficit in hippocampal slices with PDE4A5 overexpression for two weeks. Representative traces shown at time zero (black trace) and at 120 min post-TBS (colored trace) for each group (E) fEPSP slope 1 minute after theta burst stimulation (Mean ± SD relative to baseline- eGFP: 272.5 ± 92.34, PDE4A5: 173 ± 42.08; N=6 animals per group, *P* = 0.0473 by Welch’s two-tailed t-test- eGFP vs. 4A5: *t* =2.403, df = 6.991) (F) deficit in maintenance of TBS-LTP as shown by average fESP slopes over the last 20 minutes (Mean ± SD of last 20 min, 100-120 min after TBS relative to baseline- eGFP: 188.3 ± 52.73, PDE4A5: 124.1 ± 29.02; N = 6 per group, *P* = 0.0319 by Welch’s two-tailed t-test, t = 2.610, df = 7.774) (G) cAMP-PDE activity in hippocampus of mice overexpressing eGFP (or) PDE4A5 at two weeks post- surgery (Mean ± SD of cAMP-PDE activity in pmol/min/µg: eGFP: 73.62 ± 8.379, PDE4A5: 305.8 ± 84.36; P = 0.0033 by Welch’s two-tailed t-test, t = 6.116, df = 4.098).

Given that the cAMP PDE activity was comparable between overexpression of PDE4A5Δ4 and overexpression of PDE4A5 (Figure 1C), the impact of PDE4A5 overexpression on TBS-LTP is likely due to protein interactions mediated by the N-terminus of PDE4A5, which are removed from PDE4A5Δ4. Furthermore, overexpression of PDE4A5 for two weeks, which resulted in a 4-5 fold increase in cAMP PDE activity relative to eGFP overexpression (Figure 4G; Mean ± SD of cAMP-PDE activity in pmol/min/µg: eGFP: 73.62 ± 8.379, PDE4A5: 305.8 ± 84.36; P = 0.0033 by Welch’s two-tailed t-test, t = 6.116, df = 4.098) also resulted in TBS-LTP deficit (Figure 4D), despite half the overall increase in cAMP-PDE activity attained at 5 weeks overexpression with PDE4A5 and PDE4A5Δ4 (Figure 1C). PDE4A5 overexpression for 2 weeks resulted in lower post-burst potentiation of fEPSP slopes measured 1 minute after TBS (Figure 4E; Mean ± SD relative to baseline- eGFP: 272.5 ± 92.34, PDE4A5: 173 ± 42.08; N=6 animals per group, *P* = 0.0473 by Welch’s two-tailed t-test- eGFP vs. 4A5: *t* =2.403, df = 6.991) and decay of LTP to near baseline levels by 120 min after TBS (Figure 4F; Mean ± SD of last 20 min, 100-120 min after TBS relative to baseline- eGFP: 188.3 ± 52.73, PDE4A5: 124.1 ± 29.02; N = 6 per group, *P* = 0.0319 by Welch’s two-tailed t-test, t = 2.610, df = 7.774).

### CA1 targeted expression of PDE4A5 is sufficient to impair theta burst LTP

To further determine whether PDE4A5 overexpression-induced TBS-LTP deficit may be focally localized to area CA1, we overexpressed eGFP or PDE4A5 in the CA1 subregion of the hippocampal formation. PDE4A5 overexpression selectively in the CA1 subregion resulted in lower induction 1 min after TBS (Figure 5A, 5B; Mean ± SD relative to baseline- eGFP: 247.7 ± 38.7, PDE4A5: 174.1 ± 27.51; N ≥ 7 animals per group; *P* = 0.0016 by Welch’s two-tailed t-test, *t* =4.188, df = 10.69). Furthermore, PDE4A5 overexpression selectively in area CA1 resulted in the decay of the late phase of TBS-LTP to baseline levels, which is significantly different from slices expressing eGFP in CA1 (Figure 5C; Mean ± SD of last 20 min, 100-120 min after last TBS relative to baseline- eGFP: 153.5 ± 23.48, PDE4A5: 114.1 ± 24.2; N ≥ 7 animals per group; *P* = 0.0072 by Welch’s two-tailed t-test, t = 3.191, df = 12.83). In slices lacking PDE4A5 expression in the area CA1 as established by post-hoc immunohistochemistry (Figure 4G and 4H), maintenance of TBS-LTP was intact and comparable to eGFP. There were no differences between eGFP and PDE4A5 -ve groups in post-tetanic potentiation (Figure 5E; Mean ± SD relative to baseline- eGFP: 247.7 ± 38.7, N = 7 animals per group; PDE4A5 -ve: 372.3 ± 139.5, N = 3; *P* = 0.2596 by Welch’s two-tailed t-test- eGFP vs. 4A5: *t* = 1.523, df = 2.133) or in the late phase of TBS-LTP (Figure 5F; Mean ± SD of last 20 min, 100-120 min after last TBS relative to baseline- eGFP: 153.5 ± 23.48, N = 7 animals per group; PDE4A5 -ve: 166.6 ± 41.11, N = 3; *P* = 0.6463 by Welch’s two-tailed t-test, t = 0.5167, df = 2.581). These results suggest that PDE4A5 overexpression in the area CA1 is sufficient to induce deficits in TBS-LTP.

**Figure 5.**
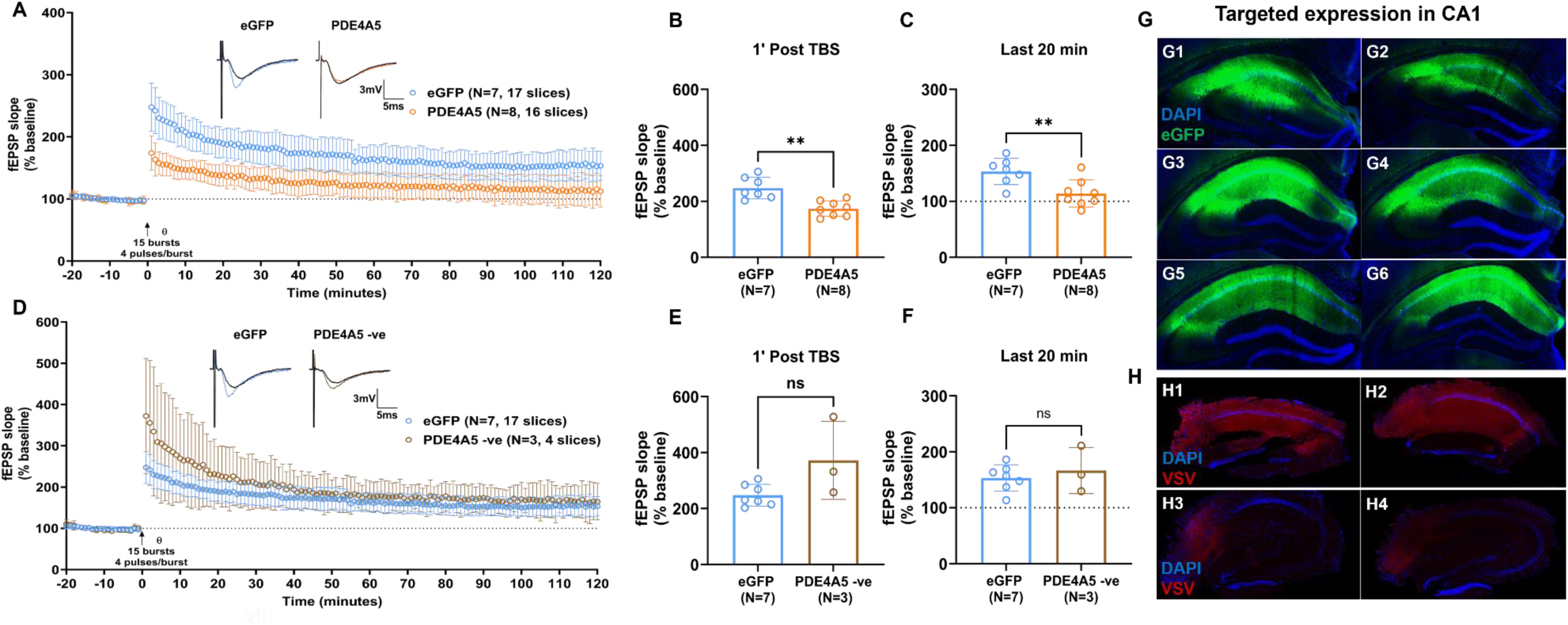
CA1 selective overexpression of PDE4A5 is sufficient to cause deficits in TBS-LTP. (A) Decay of TBS-induced LTP to baseline in slices with PDE4A5 overexpression. Representative traces are shown at time zero (black trace) and at 120 min post-TBS (colored trace) for each group. (B) fEPSP slope 1 minute after theta burst stimulation (Mean ± SD relative to baseline- eGFP: 247.7 ± 38.7, PDE4A5: 174.1 ± 27.51; N ≥ 7 animals per group; *P* = 0.0016 by Welch’s two-tailed t-test, *t* =4.188, df = 10.69) (C) deficit in the maintenance of TBS-LTP (Mean ± SD of last 20 min, 100-120 min after last TBS relative to baseline- eGFP: 153.5 ± 23.48, PDE4A5: 114.1 ± 24.2; N ≥ 7 animals per group; *P* = 0.0072 by Welch’s two-tailed t-test, t = 3.191, df = 12.83) (D) Slices lacking PDE4A5 overexpression in area CA1 as determined by post-hoc immunohistochemistry were categorized as PDE4A5 negative (PDE4A5 -ve). TBS LTP maintenance is comparable between eGFP and slices lacking PDE4A5 overexpression (PDE4A5 -ve) (E) fEPSP slope 1 minute after theta burst stimulation (Mean ± SD relative to baseline- eGFP: 247.7 ± 38.7, N = 7 animals per group; PDE4A5 -ve: 372.3 ± 139.5, N = 3; *P* = 0.2596 by Welch’s two-tailed t-test- eGFP vs. 4A5: *t* = 1.523, df = 2.133) (F) TBS-LTP maintenance is comparable between eGFP and PDE4A5 -ve (Mean ± SD of last 20 min, 100-120 min after last TBS relative to baseline- eGFP: 153.5 ± 23.48, N = 7 animals per group; PDE4A5 -ve: 166.6 ± 41.11, N = 3; *P* = 0.6463 by Welch’s two-tailed t-test, t = 0.5167, df = 2.581) (G) eGFP expression in serial sections collected 200µm apart to validate restricted expression of eGFP in area CA1 (H) representative images of slices that underwent LTP recording with (H1, H2) PDE4A5 overexpression primarily in area CA1 or (H3, H4) lack PDE4A5 overexpression in area CA1 as established using VSV-G antibody. The restricted spread of AAV for expressing PDE4A5 in area CA1 allowed for recordings in slices with or without PDE4A5 overexpression in area CA1. Slices lacking PDE4A5 overexpression in area CA1 showed low to moderate expression of PDE4A5 in the subiculum.

## Discussion

With over 25 isoforms in the PDE4 family, the underlying physiological importance of many isoforms has yet to be defined. These isoforms are generated as splice variants and through alternate promoter usage from 4 genes and, therefore, limit the utility of knockout models, making the identification of the function for each isoform a difficult task. Knockout models of each of the PDE4 subfamilies have been previously described, and subfamily functions have been identified: PDE4A or PDE4B knockout leads to increased anxiety-related behavior (Zhang *et al*., 2008; Hansen *et al*., 2014), whereas PDE4B knockouts have impaired reversal learning (Rutten *et al*., 2011). Investigation of the effects of PDE4D knockout on memory suggests distinct effects on memory depending on the behavioral task employed (Rutten *et al*., 2008; Li *et al*., 2011), but LTP enhancement in this line suggests that PDE4D subfamily isoforms may function as a brake on synaptic potentiation (Rutten *et al*., 2008). Limitations with knockout models of PDE4 subfamilies related to the lack of specificity for individual isoforms led us to study isoform-specific functions within the PDE4A subfamily by overexpression in excitatory neurons *in vivo*. We have previously reported a deficit in the consolidation of both spatial memory and contextual fear memory by overexpression of PDE4A5, the longest isoform of the PDE4A subfamily (Havekes *et al*., 2016a). This effect was isoform specific because PDE4A1, another isoform of the PDE4A subfamily, did not affect memory consolidation. In this study, we have extended our work on PDE4A subfamily isoforms to investigate hippocampal synaptic plasticity, which is believed to underlie memory consolidation (Martin *et al*., 2000; Takeuchi *et al*., 2014).

In line with our studies on memory consolidation (Havekes *et al*., 2016a), overexpression of PDE4A5, but not PDE4A1, caused impairments in TBS-LTP, although neither full-length PDE4A5 nor PDE4A1 interfered with long-lasting LTP induced by spaced 4-train stimulation. Importantly, the N-terminus of PDE4A5 was essential to mediate the effect on TBS-LTP because the lack of N-terminus in the same construct left TBS-LTP intact. Together, these results suggest that PDE4A5 overexpression results in a specific effect on LTP induction and maintenance such that some but not all forms of cAMP/PKA-dependent LTP are affected. Given that the TBS deficit is selective to PDE4A5 and requires an intact N-terminus for this deficit, our results also suggest that PDE4A5 through its N-terminus may function in a specific subcellular domain or in signalosomes (Havekes *et al*., 2016a) that interfere with TBS induced LTP, but not with L-LTP induced by spaced 4-train stimulation. Additionally, since TBS-LTP is believed to involve feed-forward inhibition of GABAergic input to the pyramidal cells (Larson & Munkácsy, 2015), an interesting case to consider in the future would be to investigate the effect of overexpression of PDE4A isoforms in excitatory cells on feed-forward inhibition.

Compartmentalization of cAMP signaling was proposed nearly half a century ago (Hayes *et al*., 1979; Hayes *et al*., 1980), and evidence in support of this compartmentalization has been provided by the identification of scaffold proteins for enzymes linked to PKA and cAMP signaling (Baillie, 2009; Lefkimmiatis & Zaccolo, 2014; Torres-Quesada *et al*., 2017; Johnstone *et al*., 2018). Although the existence of scaffold proteins has primarily supported a role for PKA compartmentalization, the evidence supporting the existence of cAMP microdomains and nanodomains in neurons is only beginning to emerge. The highly diffusible nature of cAMP, together with low V_max_ for most phosphodiesterase enzymes, challenged the notion of compartmentalized cAMP (Bock *et al*., 2020; Zhang *et al*., 2020). Recent advancements in imaging, together with the development of sensitive probes, helped identify that most cAMP exists in a buffered state in the cells and that phosphodiesterase enzymes shape these domains by degradation of cAMP in the immediate vicinity, restricting the cAMP signals to remain at baseline levels in the bulk of cytosol (Bock *et al*., 2020; Zhang *et al*., 2020). Interestingly, the sphere of influence of phosphodiesterases determined using cAMP nanorulers appears to be non-uniform, with the radius of cAMP degradation being different for different families of phosphodiesterases (Bock *et al*., 2020). Whether this property is different among isoforms of the same subfamily of phosphodiesterases needs further investigation. Such follow up investigations may provide insights into the selective deficit in TBS-LTP and memory caused by PDE4A5 overexpression but not the PDE4A1 isoform or PDE4A5Δ4.

Furthermore, compartmentalized cAMP and PKA signaling have been suggested to be critical for neurite outgrowth and axon guidance (Boczek *et al*., 2019; Boczek *et al*., 2021). In these studies, PDE4D3 anchored by AKAP6 was shown to play a pivotal role in maintaining cAMP pools in the perinuclear region, allowing for increases in cAMP levels and PKA activity as well as neurite extension (Boczek *et al*., 2019; Boczek *et al*., 2021). Whether such interactions and compartmentalized cAMP pools also facilitate cellular processes underlying memory and synaptic plasticity are not completely understood, and future work may provide insights that could explain the deficits in TBS-LTP caused by overexpression of PDE4A5 as well as identify proteins that interact with its unique N-terminus.

Regarding our findings of a selective deficit induced by overexpression of PDE4A5 on certain forms of cAMP-dependent LTP, the impact of PDE4A isoform on LTP may depend in part on the microdomains of cAMP that are targeted. Such a pattern-specific effect on cAMP forms of LTP has been previously described in the R(AB) mice, which have an attenuated PKA activity (Abel *et al*., 1997). In the R(AB) mice, where a dominant negative form of the RIα subunit of PKA is overexpressed, only LTP induced by space 4-train protocol (Abel *et al*., 1997; Woo *et al*., 2000) is affected, while LTP induced by TBS is spared (Woo *et al*., 2000). The existence of RII subunits as anchored forms enriched in dendrites (Zhong *et al*., 2009) may suggest an essential role for RII containing PKA in TBS-LTP (Woo *et al*., 2000).

The literature provides additional insight into why PDE4A5 overexpression might selectively impair TBS-LTP. The dependence of repeated spaced tetanization-induced LTP and TBS-LTP on cAMP/PKA signaling is well established (Frey *et al*., 1993; Huang & Kandel, 1994; Abel *et al*., 1997; Nguyen & Kandel, 1997; Woo *et al*., 2003). Our previous findings comparing the effects of spaced and massed tetanization on cAMP and PKA suggest that spaced tetanization allows for temporal summation of cAMP/PKA transients and results in a significantly higher increase in overall PKA activation (Kim *et al*., 2010). Perhaps, repeated tetanization in the spaced 4-train paradigm may overcome the cAMP threshold needed for expression of long-lasting LTP despite the increased cAMP PDE activity due to PDE4A5 overexpression (Kim *et al*., 2010). Because the induction of TBS-LTP involves a single train of burst stimulation that is delivered within a few seconds (3 seconds in our experiments), no further summation of cAMP/PKA transients are possible to enable the expression of long-lasting LTP. Future experiments will test the effects of PDE4A5 overexpression on LTP induced by multiple trains of TBS.

Brain-derived neurotrophic factor (BDNF), an important mediator of synaptic plasticity is released in response to neuronal activity (Lu, 2003; Lu *et al*., 2008), and PKA inhibition impairs the secretion of BDNF in cultured hippocampal neurons (Kolarow *et al*., 2007). Mice with reduced BDNF levels show intact spaced 4-train LTP (Patterson *et al*., 2001; Barco *et al*., 2005), but impaired TBS-LTP (Patterson *et al*., 2001; Barco *et al*., 2005) and synaptic tagging (Barco *et al*., 2005). Given the parallel between the selective impairment of TBS-LTP by PDE4A5 and BDNF mutants (Patterson *et al*., 2001), PDE4A5 may function upstream of BDNF signaling by gating the release and or processing of this critical neurotrophin in an activity-dependent fashion. An interesting future experiment would be to investigate the effects of PDE4A5 overexpression on synaptic tagging. A deficit in synaptic tagging due to PDE4A5 overexpression as seen in the BDNF mutants would imply that PDE4A5 regulates the localized effects on cAMP gradients to shape activity dependent BDNF release and signaling. This potential link between PDE4A5 and activity induced BDNF release may help explain the selective deficit in TBS-LTP due to PDE4A5 overexpression.

In summary, our data highlight the need to investigate the isoform-specific effects of PDE4 subfamilies and other PDE families on synaptic plasticity to enable the development of effective therapies for disorders that impair cognitive function and may benefit from reinstatement of cAMP signaling pathways.

## Data availability statement

All the contributions presented in the study are included in the data presented in the article. Any requests for data can be directed to the corresponding author.

## Author Contributions

S.M.T., T.A. conceived and designed research; S.M.T., E.N.W., and U.M. performed experiments; S.M.T., E.N.W., and U.M. analyzed data; S.M.T. and T.A. interpreted results of experiments; S.M.T. prepared figures; S.M.T. and T.A. drafted manuscript; S.M.T., and T.A. edited and revised manuscript; S.M.T., E.N.W., U.M., and T.A. approved final version of the manuscript.

## Acknowledgements

We thank Dr. Mahesh Shivarama Shetty for his guidance on electrophysiology rig setup and recordings. We also thank Dr. Yong-Seok Lee, Dr. Susan Patterson, Benjamin Kelvington and Pravda Quinoñes for their valuable feedback on this manuscript. We acknowledge the services provided by the Neural Circuits and Behavior Core for surgical procedures and microscopy experiments. Work in this manuscript is supported by grant funding from the National Institutes of Health R01MH117964 and the University of Iowa Hawk Intellectual and Developmental Disability Research Center P50 HD 103556.

